# Feedback loops in the Plk4–STIL–HsSAS6 network coordinate site selection for procentriole formation

**DOI:** 10.1101/591834

**Authors:** Daisuke Takao, Koki Watanabe, Kanako Kuroki, Daiju Kitagawa

## Abstract

Centrioles are duplicated once in every cell cycle, ensuring the bipolarity of the mitotic spindle. Although the core components have been identified, how they cooperate to achieve high fidelity in centriole duplication remains poorly understood. Here, we show that in centriole duplication the accumulation of STIL and HsSAS6, components of the cartwheel structure, provides negative feedback in the centriolar dynamics of Plk4. By live-cell imaging of endogenously tagged proteins in human cells throughout the entire cell cycle, we quantitatively tracked the dynamics of the critical duplication factors: Plk4, STIL, and HsSAS6. Centriolar Plk4 peaks and then starts decreasing during the late G1 phase, which coincides with the accumulation of STIL at centrioles. Shortly thereafter, the HsSAS6 level increases steeply at the procentriole assembly site. We also show that both STIL and HsSAS6 are necessary for attenuating Plk4 levels. Furthermore, our mathematical modeling and simulation convincingly reproduce the dynamics of these three proteins at centrioles, and suggest that the STIL–HsSAS6 complex in the cartwheel has a negative feedback effect on centriolar Plk4. Combined, these findings illustrate how the dynamic behavior of and interactions between the critical duplication factors coordinate the centriole-duplication process.

## INTRODUCTION

Centrosomes and their core components, centrioles, are duplicated once during the cell cycle, which allows bipolar spindle assembly and ensures the proper proliferation of most animal cells (Nigg and Holland, 2018; Gönczy and Hatzopoulos, 2019). Despite its significance, the precise mechanisms regulating centriole duplication are largely unknown. In human cells, three centriolar proteins, namely Polo-like kinase 4 (Plk4), STIL, and HsSAS6, have been identified as the core components that coordinate the onset of centriole duplication (Banterle and Gönczy, 2017; Nigg and Holland, 2018). Observations based on immunofluorescence microscopy have revealed a sequential accumulation of these components at centrioles during the duplication process. In the early G1 phase of the cell cycle, Plk4 first appears as a biased ring-like pattern surrounding the centrioles where STIL and HsSAS6 are still absent (Kim et al., 2013; Ohta et al., 2014, 2018). It has been suggested that the intrinsic properties of Plk4 drive this initial bias, thus providing a potential site for procentriole assembly independently of the centriolar loading of STIL and HsSAS6 (Yamamoto and Kitagawa, 2018; Takao et al., 2018). Subsequently, the ring-like pattern of Plk4 changes dynamically into a single focus containing both STIL and HsSAS6 (Arquint et al., 2015; Ohta et al., 2018, 2014). This is achieved by Plk4 binding to and phosphorylating STIL/Ana2, which subsequently stimulates the kinase activity of Plk4 (Moyer et al., 2015; Dzhindzhev et al., 2017; McLamarrah et al., 2018; Ohta et al., 2018). The phosphorylation by Plk4 of the STAN motif in STIL/Ana2 promotes the formation of a complex between the phosphorylated STIL/Ana2 and HsSAS6/DSas-6, presumably leading to the cartwheel assembly (Dzhindzhev et al., 2014; Kratz et al., 2015; Moyer et al., 2015; Arquint et al., 2015; Ohta et al., 2014). Thus, complex interactions between these components cause positive and negative regulation, which mediate the local restriction of Plk4, STIL, and HsSAS6 at the procentriole assembly site to allow for procentriole formation (Ohta et al., 2018; Arquint et al., 2015; Ohta et al., 2014).

Such sequential accumulation of the components into the centrioles has been observed mainly with fixed cells. To achieve a fundamental understanding of centriole duplication processes, however, the dynamics of the components (e.g., timing, correlation, and the interdependency of centriolar accumulation) needs to be more precisely analyzed. While live imaging has been successfully used to visualize the dynamics of these components at centrioles throughout the entire cell cycle in *Drosophila* (Aydogan et al., 2018, 2019) and *Caenorhabditis elegans* (Dammermann et al., 2008) embryos, gaining insight into these dynamic processes in human cells has remained challenging. In this study, we use optimized live imaging throughout the entire cell cycle in cultured human cells to precisely analyze and describe the dynamics of endogenous proteins participating in centriole duplication. We also simulate the dynamic processes and propose a model that explains how the dynamics of these components cooperatively organize centriole duplication.

## RESULTS AND DISCUSSION

### Distinct time courses of centriolar accumulation of endogenous Plk4, STIL, and HsSAS6 during the cell cycle

To track the behavior of endogenous proteins in live cells, we observed HCT116 cell lines by spinning disc confocal microscopy, as previously described (Takao et al., 2018). Since centriole duplication is sensitive to the expression level of the core components (e.g., overexpression of a component is known to induce overduplication of centrioles), we used endogenous tagging of target proteins. However, given the limited number of copies of endogenous centriole duplication components (Bauer et al., 2016), the signal from an endogenous fluorescent tag could be too weak to detect in live imaging. To address this issue, as previously demonstrated in both *Drosophila* embryos (Aydogan et al., 2018, 2019) and cultured human cells (Takao et al., 2018), we successfully used spinning disc confocal microscopy with an EMCCD camera to track the dynamics of endogenous proteins at centrioles. This avoided significant photobleaching of the fluorescent tag and phototoxicity to the cells throughout the entire cell cycle. In addition to Plk4 (Takao et al., 2018), we also endogenously tagged STIL and HsSAS6 with fluorescent proteins at their C-termini using the CRISPR-Cas9 system with optimized C-terminal tagging vectors (Figures 1A and S6) (Natsume et al., 2016). In the live-cell imaging, Z-stacks of fluorescence images were acquired every 10 min for up to 30 h. The cells that normally lasted at least one entire cell cycle (typically around 16 h for HCT116 cells) in the total image acquisition period were used for all data analyses, to ensure that we used only cells that had entered their next cell cycle without phototoxicity.

**Figure 1.**
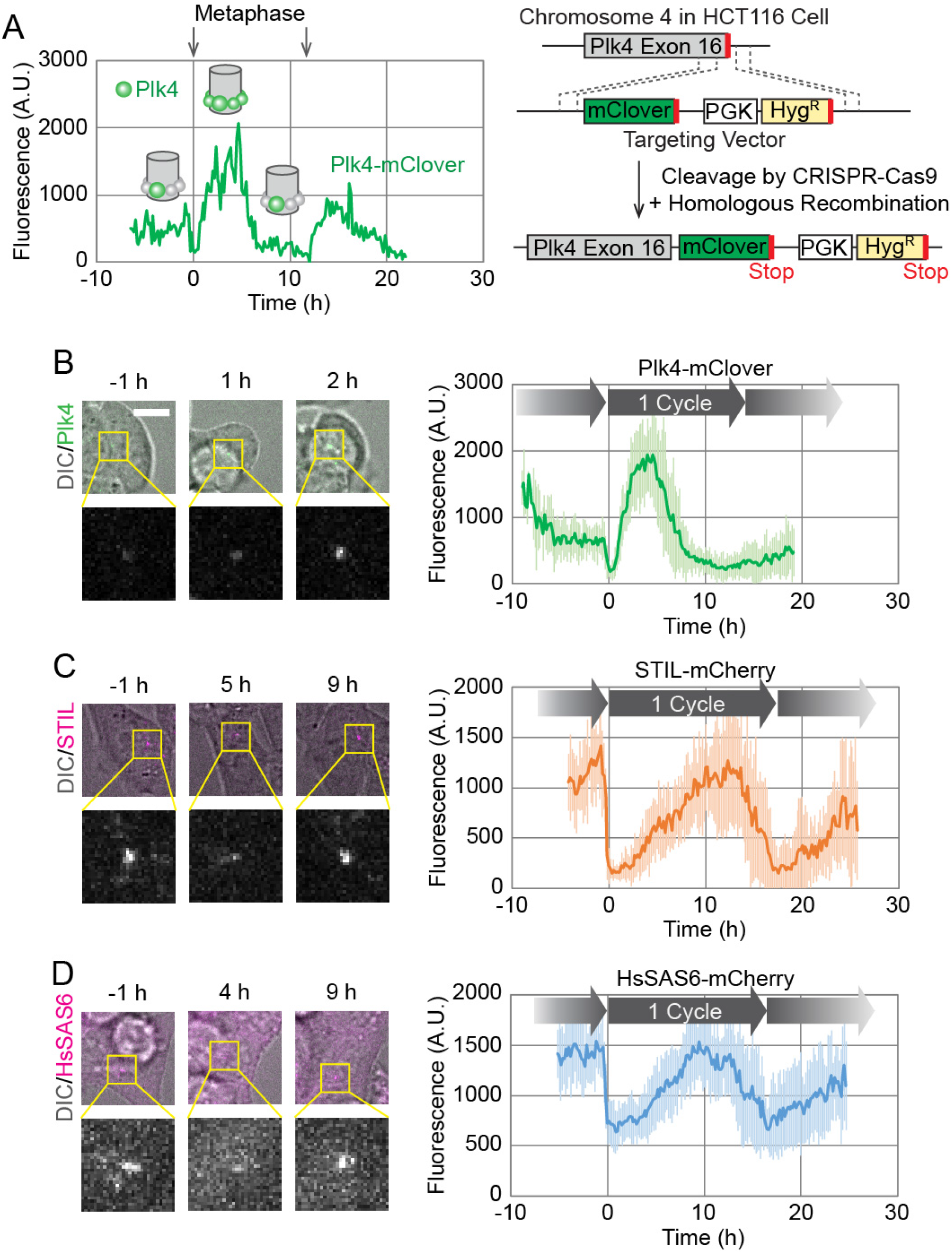
Live imaging of endogenously tagged proteins involved in centriole duplication. (A) Time course of centriolar Plk4-mClover fluorescence from a single cell. The cell divided twice during the 30 h observation period, as indicated by the arrows showing metaphase. Schematics in the graph show putative spatial patterns of Plk4 around the mother centriole at the corresponding time points. The endogenous tagging system is schematically shown on the right. (B–D) Averaged time courses of Plk4-mClover (B), STIL-mCherry (C), and HsSAS6-mCherry (D) signals at the centrioles of 14, 11, and 12 cells, respectively. Time course data were aligned at metaphase (0 h). The period between two metaphase time points is defined as one cycle. Note that the length of the cycle varies slightly among the averaged graphs due to the variety in the cell population. Error bars, SD. Representative images are shown on the left of each graph. Scale bar, 10 μm.

First, we confirmed that the fluorescence intensity of Plk4-mClover oscillated in concert with the cell cycle in human cells (Figures 1A and 1B). This oscillation has been shown to reflect the changes in the spatial pattern of centriolar Plk4, i.e., from the ring-like to the single-focus pattern (Takao et al., 2018), as schematically shown in the graph in Figure 1A. To further verify the behavior of Plk4-mClover at centrioles, we monitored the effect of treatment with a Plk4 inhibitor, centrinone (Wong et al., 2015). Centrinone treatment is known to promote centriolar accumulation of Plk4 in a few hours, presumably by inhibiting dissociation and/or degradation of Plk4 (Ohta et al., 2018). Indeed, the centriolar Plk4-mClover signal increased five-to tenfold immediately following the addition of centrinone, regardless of the stage of interphase (Figure S1), suggesting that centriolar accumulation of Plk4 is tightly regulated by its phosphorylation state during interphase. Interestingly, during mitosis, the Plk4-mClover signal decreased, even in the presence of centrinone (Figure S1).

We then similarly observed the dynamics of STIL-mCherry and HsSAS6-mCherry at centrioles throughout the cell cycle (Figures 1C and 1D). In contrast to Plk4-mClover, which started increasing immediately after mitotic exit (Figure 1B), the fluorescence intensity of STIL-mCherry and HsSAS6-mCherry started increasing about 2–5 h after mitotic exit, and decreased just before the next mitosis (Figures 1C and 1D). This is consistent with previous observations, in which endogenous or exogenously-expressed STIL and HsSAS6 were absent from centrioles during the early G1 phase (Strnad et al., 2007; Arquint et al., 2012; Arquint and Nigg, 2014). To further investigate whether STIL and HsSAS6 are indeed absent from centrioles during the postmitotic time window, we attempted to detect signals with higher sensitivity by switching tags from mCherry to mClover, one of the brightest fluorescent proteins. Both time courses were quite similar (Figures S2A and S2B), but on closer inspection a weak centriolar signal of HsSAS6-mClover was detectable during and immediately after mitosis (Figure S2C). It is surprising that HsSAS6 is already present around centrioles during the early G1 phase, albeit at very low levels. This discrepancy from previous observations may stem from differences in the immunofluorescence protocols. One possibility is that the centriolar fraction of HsSAS6 during the early G1 phase is not in a rigid structure like cartwheels, so these molecules would not be retained at centrioles during the fixation procedure. This observation aside, the majority of the centriolar HsSAS6 we observed started accumulating at the centrioles later in the cell cycle, regardless of the tags we used (Figures 1D and S2B), as previously observed via immunofluorescence (Keller et al., 2014; Strnad et al., 2007). This centriolar HsSAS6 thus probably corresponds to the stack of cartwheels, at least in part. The meaning of the fraction of centriolar HsSAS6 that is present during the early G1 phase is yet to be addressed. It is possible that it provides the seed for the onset of cartwheel assembly. Alternatively, it could merely be a meaningless fraction that happens to be trapped in the crowded pericentriolar environment (Fu and Glover, 2012; Lawo et al., 2012; Mennella et al., 2012; Sonnen et al., 2012; Woodruff et al., 2017). Regardless, this approach of optimized live-cell imaging with endogenous tagging illustrates the distinct dynamics of critical centriole duplication factors throughout the cell cycle in human cells.

### Centriolar accumulation of STIL coincides with the onset of the drop in Plk4, shortly followed by the steep increase in centriolar HsSAS6

We then sought to investigate how the centriolar dynamics of the core duplication components correlate with each other. To this end, we generated cell lines in which both Plk4 and STIL or HsSAS6 were endogenously tagged with mClover and mCherry, respectively. Data from simultaneous live imaging of the two proteins are shown in Figures 2A, 2B, S3, and S4. To precisely compare the time courses of the two proteins, data were not pooled as in Figures 1B–D; rather, data from single cells are shown individually (smoothed by applying a moving average through time for noise reduction). In contrast to the averaged curves, both STIL- and HsSAS6-mCherry tended to exhibit a steep and sometimes stepwise increase in fluorescence upon centriolar accumulation (Figures 2A, 2B, S3, and S4).

**Figure 2.**
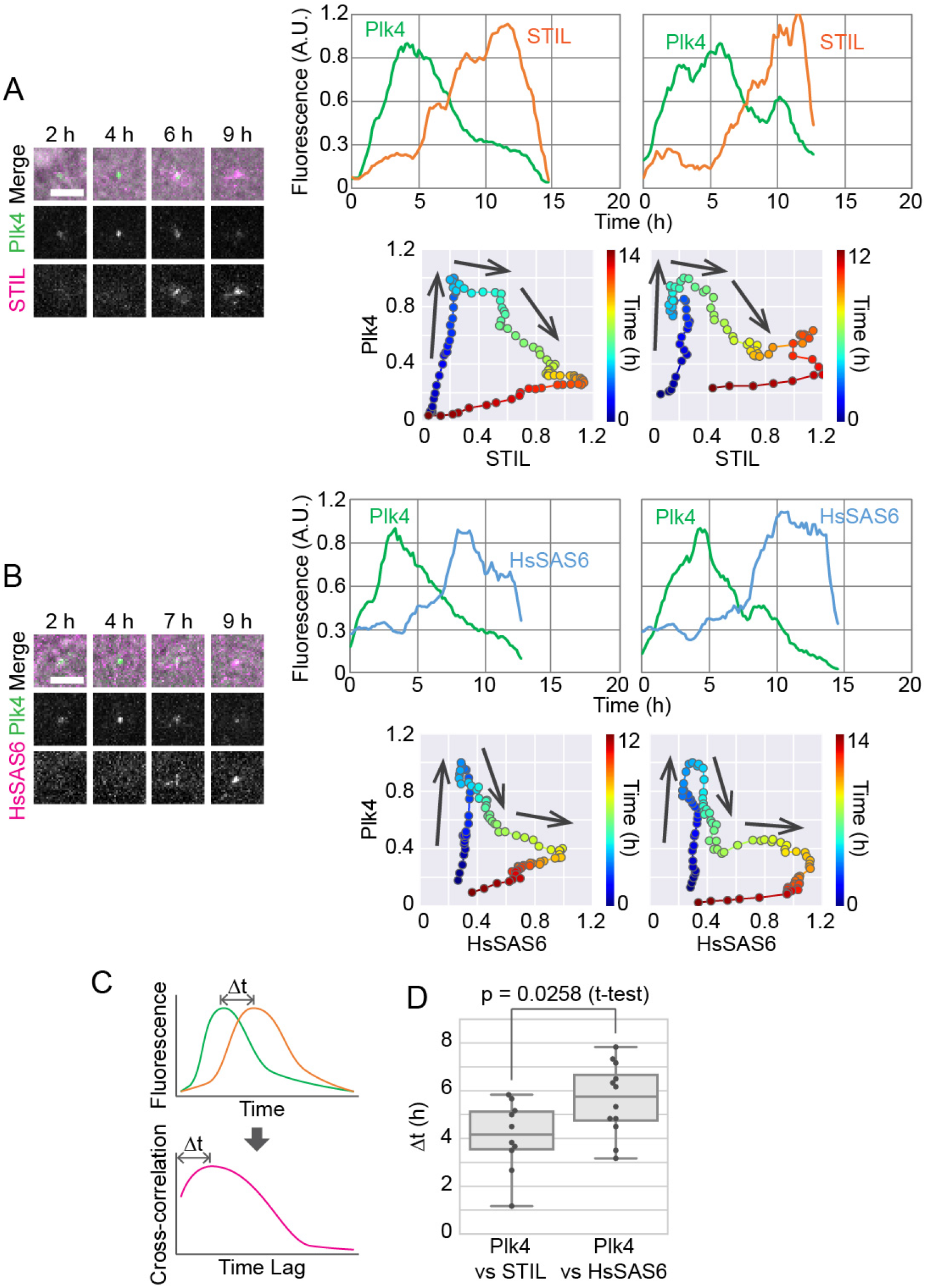
Distinct behavior of STIL and HsSAS6 relative to Plk4 in centriolar accumulation. (A, B) Simultaneous imaging of endogenously tagged Plk4 and STIL (A) or HsSAS6 (B). Representative images (left) and two example graphs (right) are shown for each. Scale bar, 5 μm. Each graph is of data from a single cell with moving-average smoothing (± 3 time points). The time course of normalized fluorescence is shown in two different ways: fluorescence of each component against time (top) and against each other with time color-coded (bottom). Arrows on the graphs indicate the direction of the time course. See also Figures S3 and S4. (C, D) Cross-correlation analysis to compare the time courses of STIL-mCherry or HsSAS6-mCherry with that of Plk4-mClover. Schematics of how to obtain the time lag (Δt) based on cross-correlation (C) and obtained Δt values (D) are shown.

The centriolar STIL-mCherry signal tended to start increasing at the peak in the time course of the centriolar Plk4-mClover signal. In other words, the centriolar Plk4-mClover signal started increasing after mitotic exit, and decreased just after the centriolar STIL-mCherry signal started increasing (Figures 2A and S3). Thus, centriolar accumulation of STIL probably triggers the decrease in centriolar Plk4. This is consistent with previous immunofluorescence observations, as well as the model in which centriolar loading of STIL promotes disassembly of the ring-like pattern of Plk4 to form the single-focus pattern (Arquint et al., 2015; Ohta et al., 2018, 2014). The centriolar HsSAS6-mCherry signal, on the other hand, only began to increase dramatically after the Plk4-mClover signal had started decreasing (Figures 2B and S4).

To further verify the sequential accumulation of STIL and HsSAS6, we next used cross-correlation analysis to compare the time lags in the centriolar accumulation of STIL-mCherry and HsSAS6-mCherry relative to Plk4-mClover (Figures 2C and 2D). For Plk4-mClover and STIL-mCherry, this time lag was on average 4.1 h, and for Plk4-mClover and HsSAS6-mCherry, it was on average 5.6 h (Figure 2D). It is therefore likely that after a certain amount of STIL has accumulated at a Plk4 focus around centrioles, HsSAS6 begins stacking cartwheels on the focus. The centriolar STIL level, similarly to that of HsSAS6, kept increasing after the drop in centriolar Plk4, suggesting that STIL and HsSAS6 are coordinately incorporated into the stack of cartwheels.

Combined, these results suggest the following: (1) elevated loading of STIL triggers the pattern shift of centriolar Plk4 from the ring-like to the single-focus pattern; (2) following the gradual accumulation of phosphorylated STIL, and once the local concentration of HsSAS6 at the centrioles exceeds the threshold for cartwheel formation, the STIL–HsSAS6 complex is coordinately integrated into a stack of cartwheels; and (3) this transition leads to a stable and continuous reduction in Plk4 at the centrioles, which may contribute significantly to the formation of a single procentriole site. STIL may thus play multiple roles as a hub in these sequential processes during centriole duplication.

### Both STIL and HsSAS6 are required, in an interdependent manner, for the spatial-pattern shift of centriolar Plk4

Since STIL and HsSAS6 are thought to induce the spatial-pattern shift of centriolar Plk4 (Arquint et al., 2015; Ohta et al., 2018, 2014), we then observed the behavior of centriolar Plk4 in cells depleted of STIL or HsSAS6 (as well as in a control culture). In live-cell imaging of endogenous Plk4-mClover after mitotic exit, the onset of centriolar accumulation and the time until it reached its peak fluorescence intensity were similar in all three conditions, although the accumulation rates seemed slightly faster in siSTIL- or siHsSAS6-treated cells than in the control (Figure 3A). In stark contrast, upon depletion of STIL or HsSAS6, centriolar Plk4-mClover remained constant after reaching its fluorescence peak throughout the rest of the period until the next mitosis (Figure 3A). The peak intensity of centriolar Plk4-mClover fluorescence in these conditions was comparable to that in the control (Figure 3A). These results suggest that, in STIL- or HsSAS6-depleted cells, Plk4 forms the centriolar ring after mitotic exit as normal, but the pattern shift to a single focus never takes place. Indeed, when observed by immunofluorescence microscopy at a higher resolution, in line with previous observations (Ohta et al., 2014), centriolar Plk4 primarily exhibited its ring-like pattern in STIL- or HsSAS6-depleted cells (Figure 3B).

**Figure 3.**
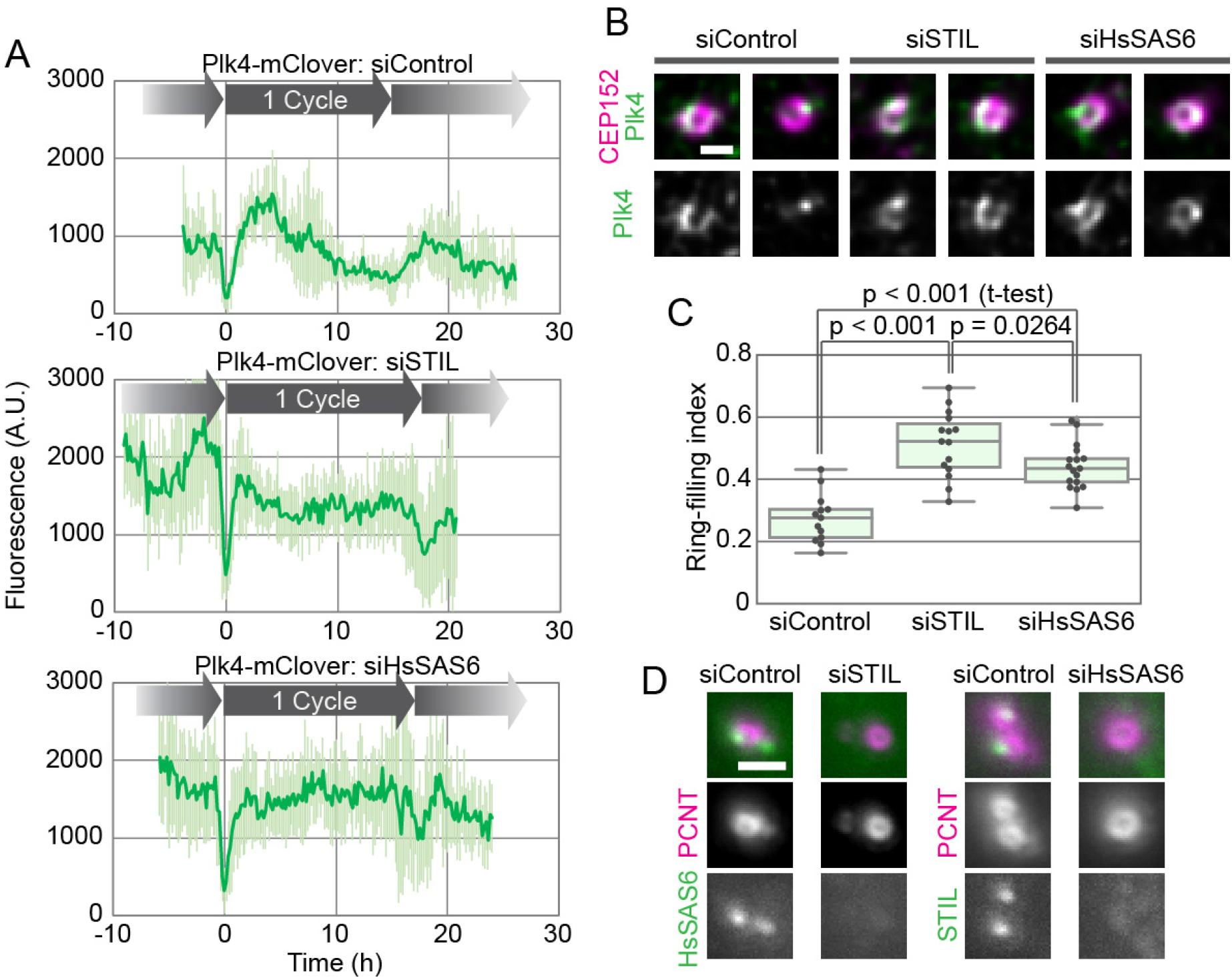
Effect of depletion of STIL or HsSAS6 on the behavior of centriolar Plk4. (A) Time course of centriolar Plk4-mClover fluorescence in cells transfected with siControl (top), siSTIL (mid), or siHsSAS6 (bottom). The results in the graphs are shown as an average ± SD of 8, 10, and 8 cells, respectively. (B, C) Effect of depletion of STIL or HsSAS6 on the spatial patterning of centriolar Plk4. Representative high-resolution immunofluorescence images of Plk4 with and without the centriole marker CEP152 (B) and the ring-filling indices (C) for the three conditions. Scale bar, 0.5 μm. (D) Interdependency of centriolar accumulation of STIL and HsSAS6. HeLa cells were transfected with siControl, siSTIL, or siHsSAS6 and then stained with antibodies to STIL or HsSAS6 and the centriole marker pericentrin (PCNT). Representative images are shown for each combination. Scale bar, 1 μm.

To quantify the spatial pattern of Plk4, we calculated ring-filling indices for centriolar Plk4 from the immunofluorescence images. The ring-filling index is a parameter defined such that the more like a ring the spatial pattern of centriolar Plk4 is, the closer to 1 the index is (Takao et al., 2018). Ring-filling indices of centriolar Plk4 in STIL- or HsSAS6-depleted cells were significantly higher than those in control cells (Figure 3C). Taken together, these data show that both STIL and HsSAS6 are required for the pattern shift of centriolar Plk4 to a single focus but are dispensable for the initial formation of the ring-like pattern of Plk4.

Centriolar accumulation of STIL and HsSAS6 is known to be interdependent (Tang et al., 2011; Arquint et al., 2012; Vulprecht et al., 2012). We confirmed, by using immunofluorescence microscopy, that STIL is mostly absent from centrioles in cells that are depleted of HsSAS6 and vice versa (Figure 3D). Therefore, STIL and HsSAS6 may cooperatively stabilize the centriolar Plk4 focus, with slightly different time courses, to provide a single site for procentriole formation. Interestingly, the ring-filling indices for Plk4 differed slightly between siSTIL- and siHsSAS6-treated cells (Figure 3C). This may reflect differences in the roles of STIL and HsSAS6 in the spatial regulation of Plk4 at centrioles, although further investigation is required.

### Centriolar accumulation of STIL and HsSAS6 may regulate the dynamics of centriolar Plk4 to generate a single site for procentriole formation

To understand the molecular mechanisms underlying procentriole formation with respect to the dynamic behavior of the participating components, we constructed a mathematical model to reproduce the experimental results by simulation (Figure 4A). Plk4 possesses intrinsic self-organization properties, such as self-assembly and the promotion of dissociation/degradation (hereafter collectively referred to as dissociation) of Plk4 molecules in an autophosphorylation-dependent manner (Yamamoto and Kitagawa, 2018). Based on these properties, as in the previous model simulating the spatial patterning of centriolar Plk4 (Takao et al., 2018), the present model assumes that the autophosphorylation-mediated activation of Plk4 promotes its dissociation from the centriole. STIL promotes the kinase activity of Plk4 and thus the dissociation of centriolar Plk4. STIL also attenuates the dissociation of phosphorylated Plk4 via direct binding. This complex regulation is thought to be mediated by the bimodal binding of STIL to Plk4 (Ohta et al., 2018). Indeed, centriolar loading of STIL is mediated by Plk4 in a phosphorylation-dependent manner. Given that HsSAS6 can be detected at centrioles during the early G1 phase (Figure S2C), centrioles can recruit HsSAS6 just after mitotic exit, but may not be able to retain much of it at that stage. Since the steep increase in centriolar STIL preceded that of HsSAS6 and continued in concert with it (Figures 2, S3, and S4), the model assumes that phosphorylated STIL mediates the cartwheel assembly via direct interaction with HsSAS6. In addition, it may be reasonable to assume that HsSAS6 is retained at centrioles once it has formed a closed ring and been incorporated into the stable cartwheel structure, resulting in a drastic increase in the centriolar HsSAS6 level. The formation of cartwheel structures may also require phosphorylated STIL as part of the stable structure, thus preventing it from dissociating. Via the interaction network described above, in our model Plk4, STIL, and HsSAS6 cooperatively generate a single site for procentriole formation.

**Figure 4.**
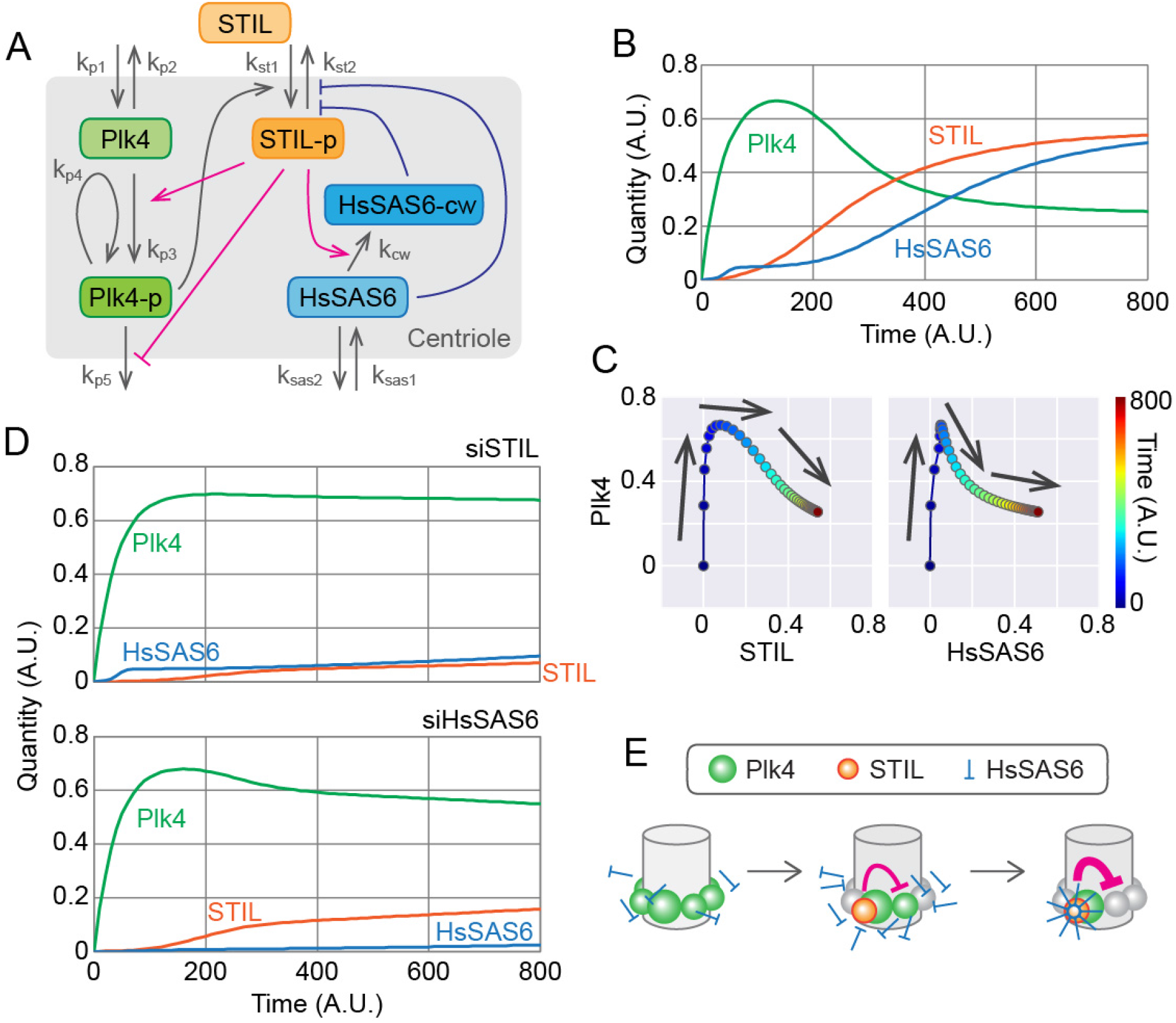
Mathematical modeling of the centriolar dynamics of Plk4, STIL, and HsSAS6. (A) Schematic of the mathematical model. Plk4-p, STIL-p, and HsSAS6-cw denote the phosphorylated forms of Plk4 and STIL, and HsSAS6 in the cartwheel structure, respectively. See the text and Methods for details. (B) Simulated time course of Plk4, STIL, and HsSAS6 at a centriole. Total quantities (Plk4 + Plk4-p, STIL + STIL-p, and HsSAS6 + HsSAS6-cw) are shown. Note that quantity and time are all relative (dimensionless). (C) Changes in the quantity of Plk4 against that of STIL or HsSAS6. The simulation data have been plotted in the same way as in the graphs in Figure 2. (D) Simulated effect of STIL or HsSAS6 depletion. Simulation results with the expression levels of STIL or HsSAS6 decreased to 10% and 1% of the normal levels, respectively, are shown. (E) Schematic of the model of the onset time course of procentriole formation.

Using the model, we simulated the time course of the molecules throughout the cell cycle. For simplicity, we have displayed the total quantity of each component at the centriole in Figure 4B, although the actual simulation included subclasses of components (e.g., the non-phosphorylated and phosphorylated forms of Plk4; Figure S5). Our live-cell imaging did not distinguish between such subclasses, so it is the data shown in Figure 4B that should be compared with the live-cell imaging data. The simulated time courses were quite similar to those in our actual observations, and the parametric plots (Plk4 vs STIL or HsSAS6) from the simulation (Figure 4C) also resembled the experimental data (Figures 2A, 2B, S3, and S4). The model could therefore convincingly reproduce the experimental data obtained under normal physiological conditions.

We then compared the simulation data with the experimental data obtained under STIL- or HsSAS6-depleted conditions. In both cases, the increased level of Plk4 was maintained over time after it had reached its peak, and the onset of Plk4 accumulation was unaffected (Figure 4D). Centriolar accumulation of HsSAS6 was significantly attenuated with the depletion of STIL, and vice versa (Figure 4D). Thus the simulations based on our model also convincingly reproduced the experimental data obtained under perturbed conditions.

As has been previously proposed (Arquint et al., 2015; Ohta et al., 2018, 2014), the spatial-pattern shift of Plk4 from the pericentriolar ring to a single focus is mediated by STIL and HsSAS6. However, how exactly the coordinated action of the three components creates a single site for procentriole assembly remains elusive. Based on both experimental observations and simulations, we propose a model of procentriole formation as schematically illustrated in Figure 4E. Our live-cell imaging of single cells throughout the cell cycle demonstrated that the centriolar loading of both STIL and HsSAS6 is tightly associated with, and indeed required for, the rapid decrease in Plk4 after it has reached its fluorescence peak. Our mathematical modeling and simulation faithfully reproduce these processes. The significance of STIL in the spatial patterning of centriolar Plk4 has also been demonstrated using another mathematical model, which assumes different behavior and interactions of Plk4, STIL, and their phosphatases (Leda et al., 2018).

Interestingly, immediately after mitotic exit, HsSAS6 can already be recruited to centrioles, presumably in the free dimer state, despite its low levels (Figures S2C and 4E). It exhibits a high dissociation constant for the self-assembly of N-terminal head domains to form a 9-fold symmetric ring (van Breugel et al., 2011; Kitagawa et al., 2011), however, so these low levels may not be sufficient to initiate cartwheel assembly. Phosphorylated STIL promotes centriolar accumulation of HsSAS6 by forming a complex that is thought to be a crucial process in cartwheel assembly (Moyer et al., 2015; Kratz et al., 2015; Ohta et al., 2014). We thus hypothesize that once the local concentration of HsSAS6 dimers exceeds the threshold for forming a closed, rigid ring as the center of the cartwheel structure, increasing numbers of HsSAS6 dimers flow into the stream of the building stack of cartwheels. This is analogous to a previous model based on an *in vitro* reconstitution assay using *Chlamydomonas reinhardtii* SAS6 and mathematical modeling, which suggested that the cartwheel assembly mechanism was mediated by the *Chlamydomonas* protein Bld10p (Klein et al., 2016; Goldie et al., 2017; Nievergelt et al., 2018). Given that STIL is thought to be a part of the cartwheel structure (Stevens et al., 2010), this system acts as a positive feedback loop for the accumulation of the STIL–HsSAS6 complex at centrioles. Concurrently, at the procentriole assembly site, this increased quantity of the protein complex may suppress the dissociation of active Plk4 from centrioles via direct interaction, while promoting the activation and subsequent dissociation of neighboring Plk4 molecules from centrioles. In this way, it is possible that the structural integrity of assembling cartwheels is directly linked with the negative feedback loop that completely suppresses the formation of extra procentrioles. Our data and these arguments suggest that positive and negative regulation within the Plk4–STIL–HsSAS6 network ensures accurate site selection for the formation of a single procentriole.

Recent work on *Drosophila* embryos has precisely described the dynamics of centriolar proteins, including Plk4 and Sas6 (Aydogan et al., 2018, 2019). The time course of *Drosophila* Plk4 is similar to that indicated by this study, although the cell cycle is much shorter (~10–20 min) and the onset of centriolar Sas6 accumulation occurs before Plk4 reaches its peak (Aydogan et al., 2018, 2019). However, given that Plk4 appears as a ring only in late mitosis, when Sas6 begins to be recruited to centrioles, similar regulation to that indicated in our study may be occurring at the onset of centriole duplication. In line with this, there seems to be a small Plk4 peak around mitotic exit in *Drosophila* embryos (Aydogan et al., 2018, 2019). Despite the homology, the precise mechanisms regulating the rate and timing of each process may differ depending on the cell, tissue type, and species. Further research, including optical imaging and molecular analyses of the protein dynamics, structural analyses, and computational analyses, will provide insight into the mechanisms underlying the tight regulation of centriole duplication.

## ACKNOWLEDGEMENTS

We gratefully acknowledge A. Kimura for advice on mathematical modeling; R. Matsuura for conducting pilot experiments; M. Ohta, S. Yoshiba, T. Natsume, and M. Kanemaki for technical support in the generation of the HCT116 cell lines used in this study; and the members of the Kitagawa laboratory for technical support, discussion, and critical review of the manuscript. This work was supported by a Grant-in-Aid for Young Scientists A and Scientific Research C from the Ministry of Education, Science, Sports and Culture of Japan and by the Takeda Science Foundation, the Daiichi Sankyo Foundation of Life Science, the Uehara Memorial Foundation, and the Japan Prize Foundation.

## AUTHOR CONTRIBUTIONS

D.T. and D.K. designed the study and experiments. D.T., K.W., and K.K. performed the experiments. D.T. performed the simulations. D.T. and K.K. analyzed the data. D.T. and D.K. interpreted the data. D.T. and D.K. wrote the manuscript.

---

**Figure S1.**
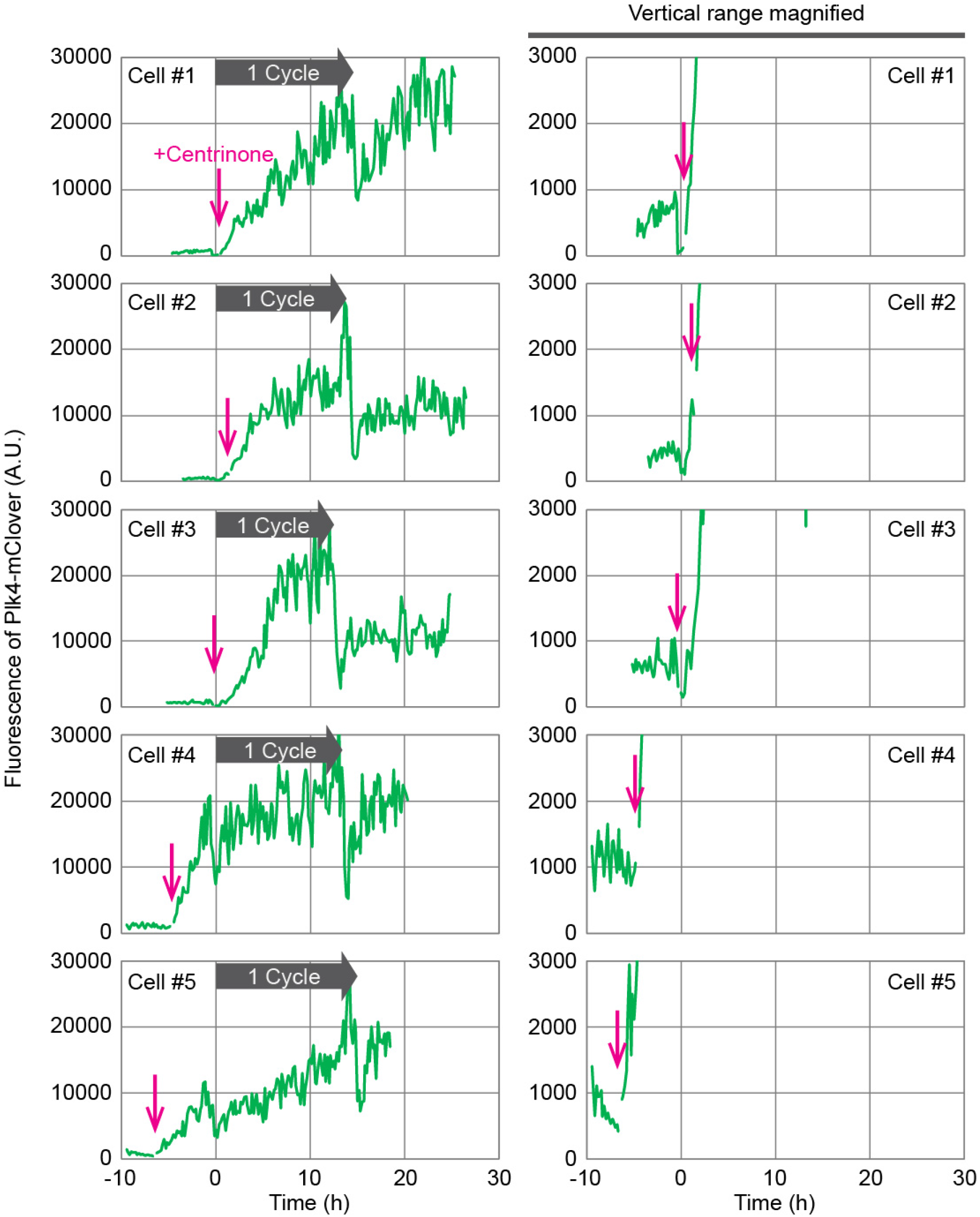
Effect of centrinone treatment on the centriolar accumulation of Plk4. Cells were treated with centrinone at various time points during the live imaging. Plk4-mClover signal was monitored as in Figure 1A, but centrinone was added to the culturing medium at the time points indicated by arrows. Representative graphs for five single cells are shown. As the fluorescence intensity of Plk4-mClover increased up to tenfold, the same data with an adjusted vertical range are shown in the righthand column, for comparison with the control data presented in Figures 1A and 1B. When the centrinone was added, image acquisition was paused for 20 min, as indicated by the gaps in the time course graphs.

**Figure S2.**
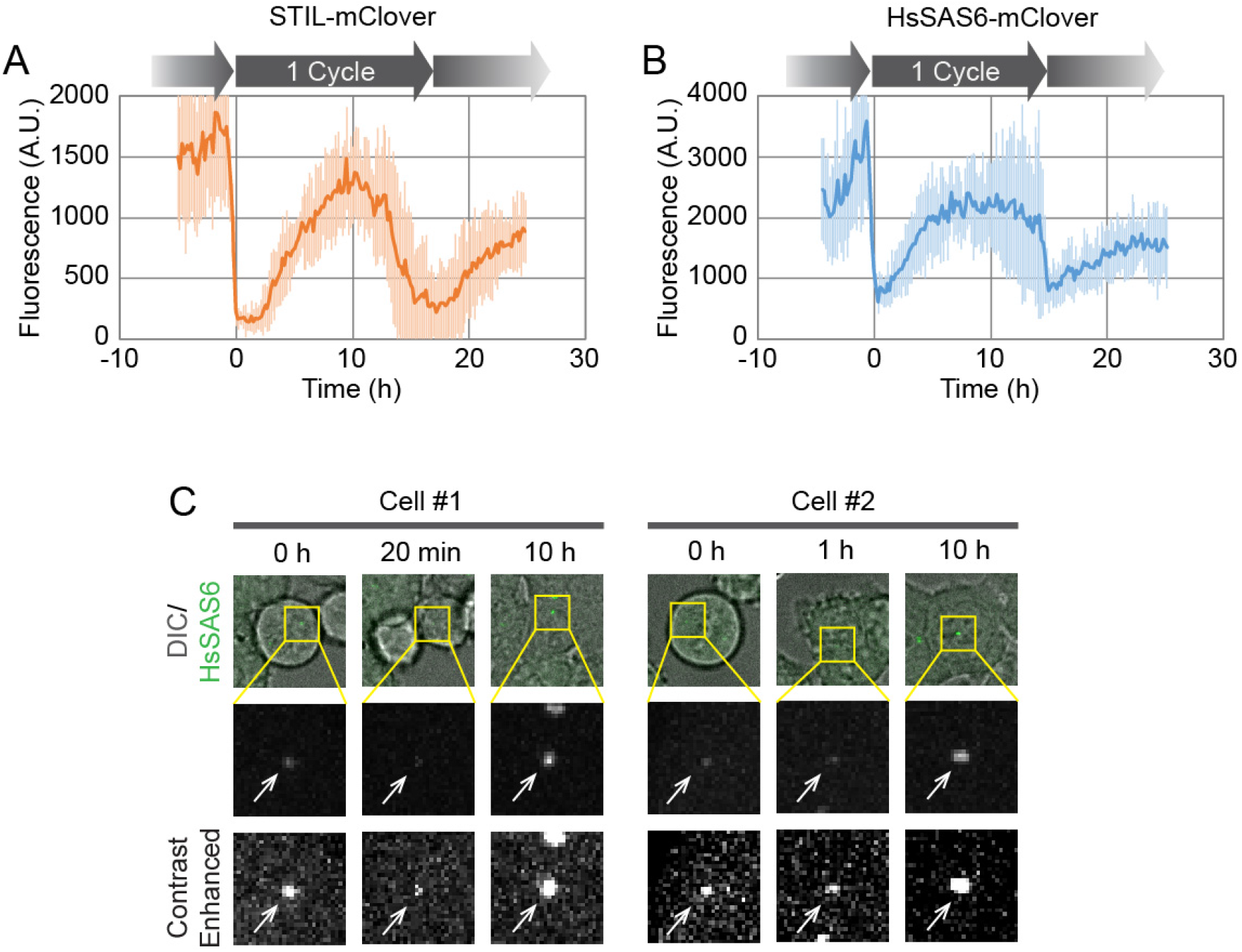
Live imaging of STIL and HsSAS6 endogenously tagged with mClover. (A, B) Average fluorescence of STIL-mClover (A) and HsSAS6-mClover (B). Data were obtained similarly to those in Figures 1C and 1D, except that the fluorescent tag used was mClover instead of mCherry. Graphs were prepared similarly to those in Figure 1. n = 11 cells each. (C) Representative images for HsSAS6-mClover from two different cells. The amount of time elapsed since metaphase is shown at the top of each column. Centriolar HsSAS6-mClover is indicated with arrows in the magnified images in the central (regular contrast) and bottom (enhanced contrast) rows.

**Figure S3.**
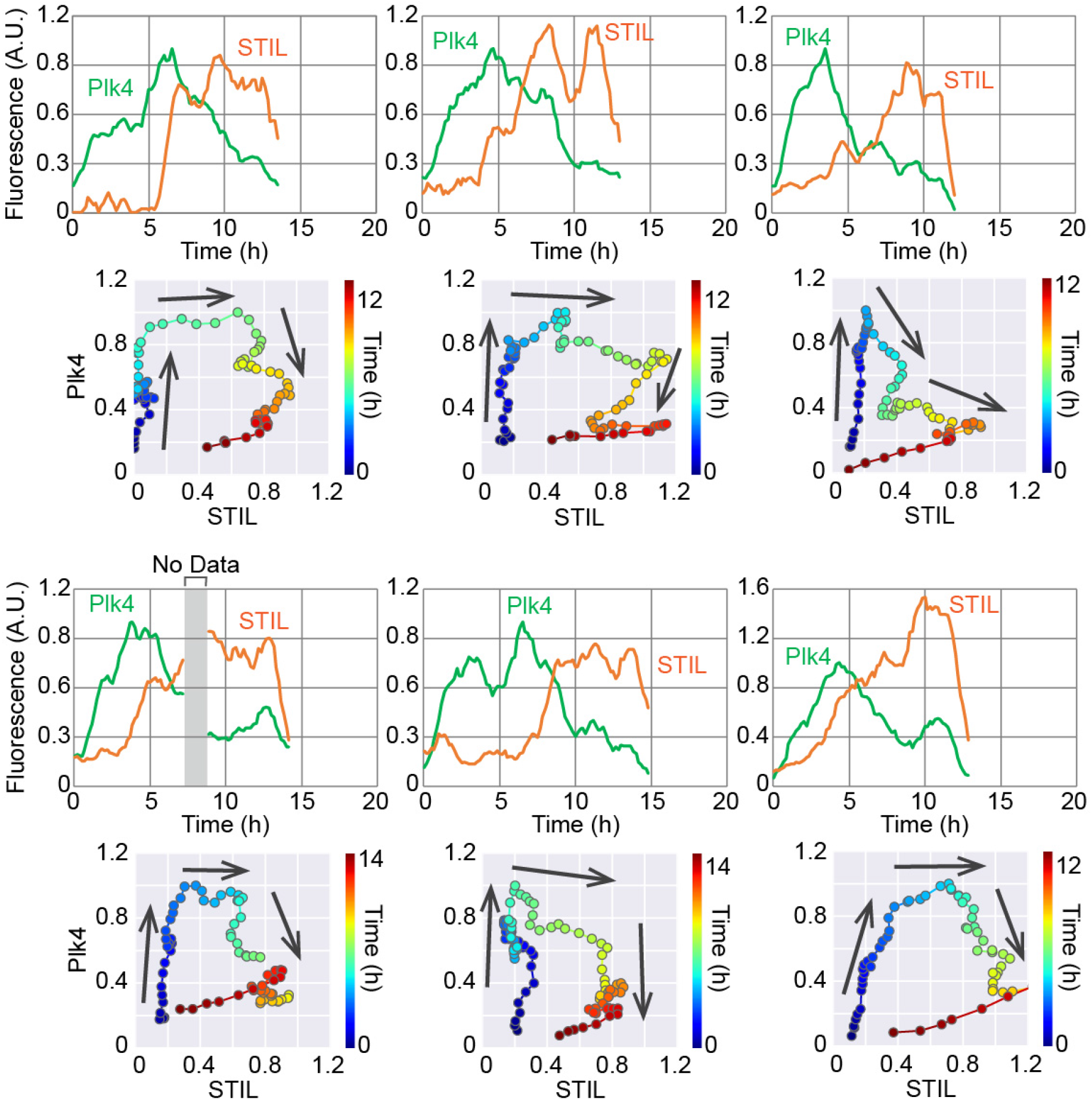
Simultaneous imaging of Plk4-mClover and STIL-mCherry. Example data supplementary to those presented in Figure 2A; the data for six additional cells are shown. The break in the graphs for one cell (shaded area labeled “No Data”) was due to an interruption in image acquisition caused by a bubble in the immersion oil.

**Figure S4.**
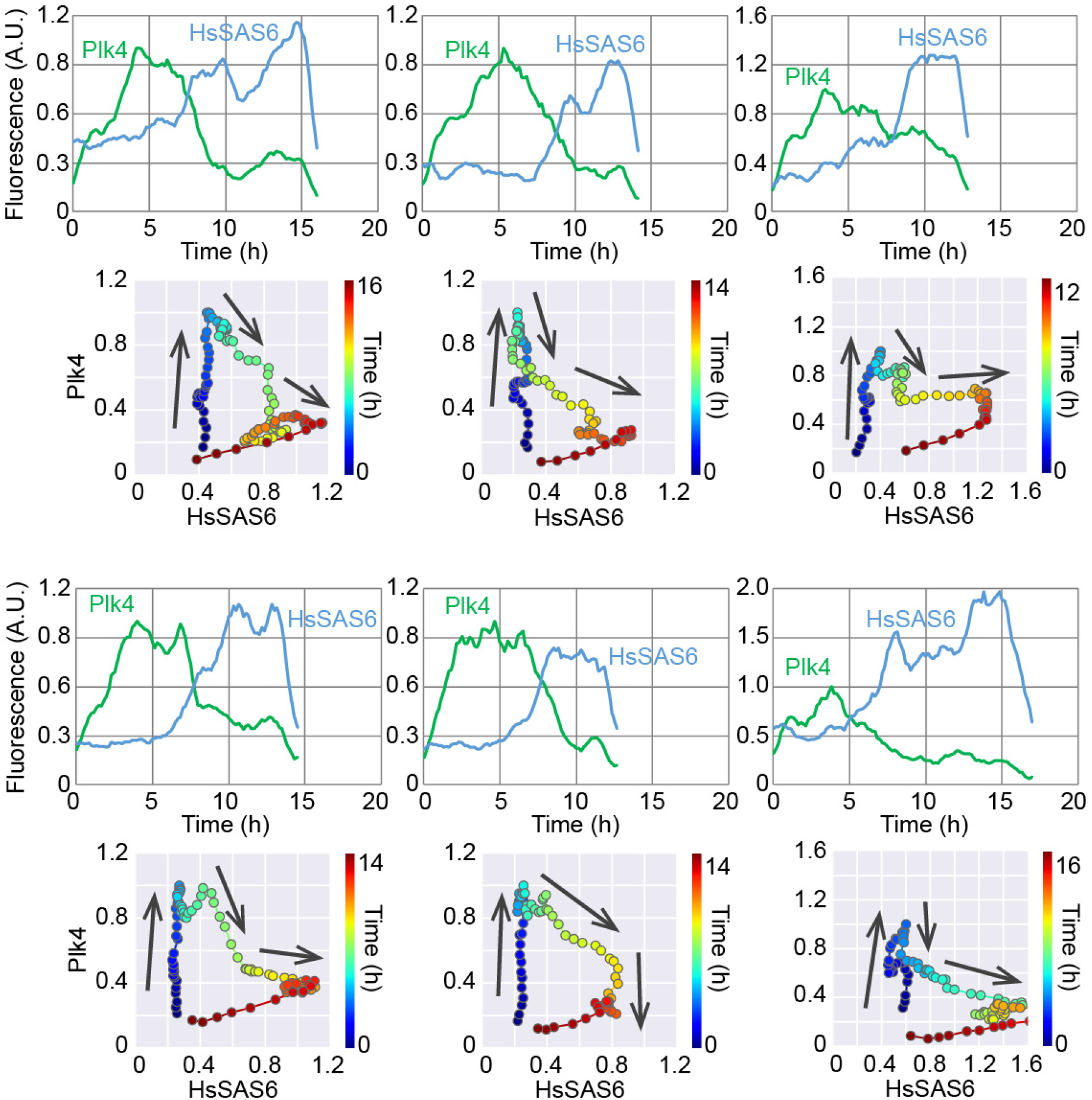
Simultaneous imaging of Plk4-mClover and HsSAS6-mCherry. Example data supplementary to those presented in Figure 2B; the data for six additional cells are shown.

**Figure S5.**
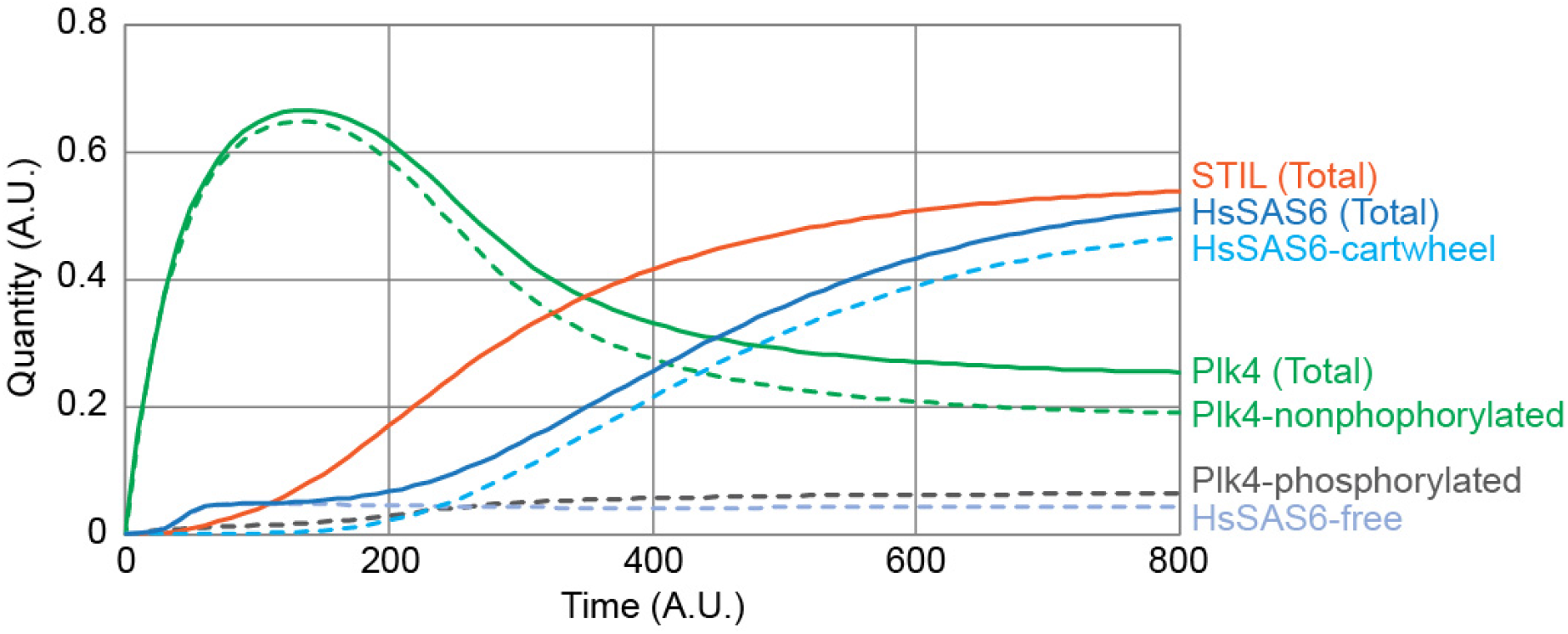
Simulation results for all the components modeled. A graph of the same data shown in Figure 4B showing the other components that were simulated but omitted from Figure 4B for simplicity. Specifically, the quantities of Plk4-non-phosphorylated, Plk4-phosphorylated, HsSAS6-free, and HsSAS6-cartwheel are shown here, in addition to the totals for each of the three proteins.

**Figure S6.**
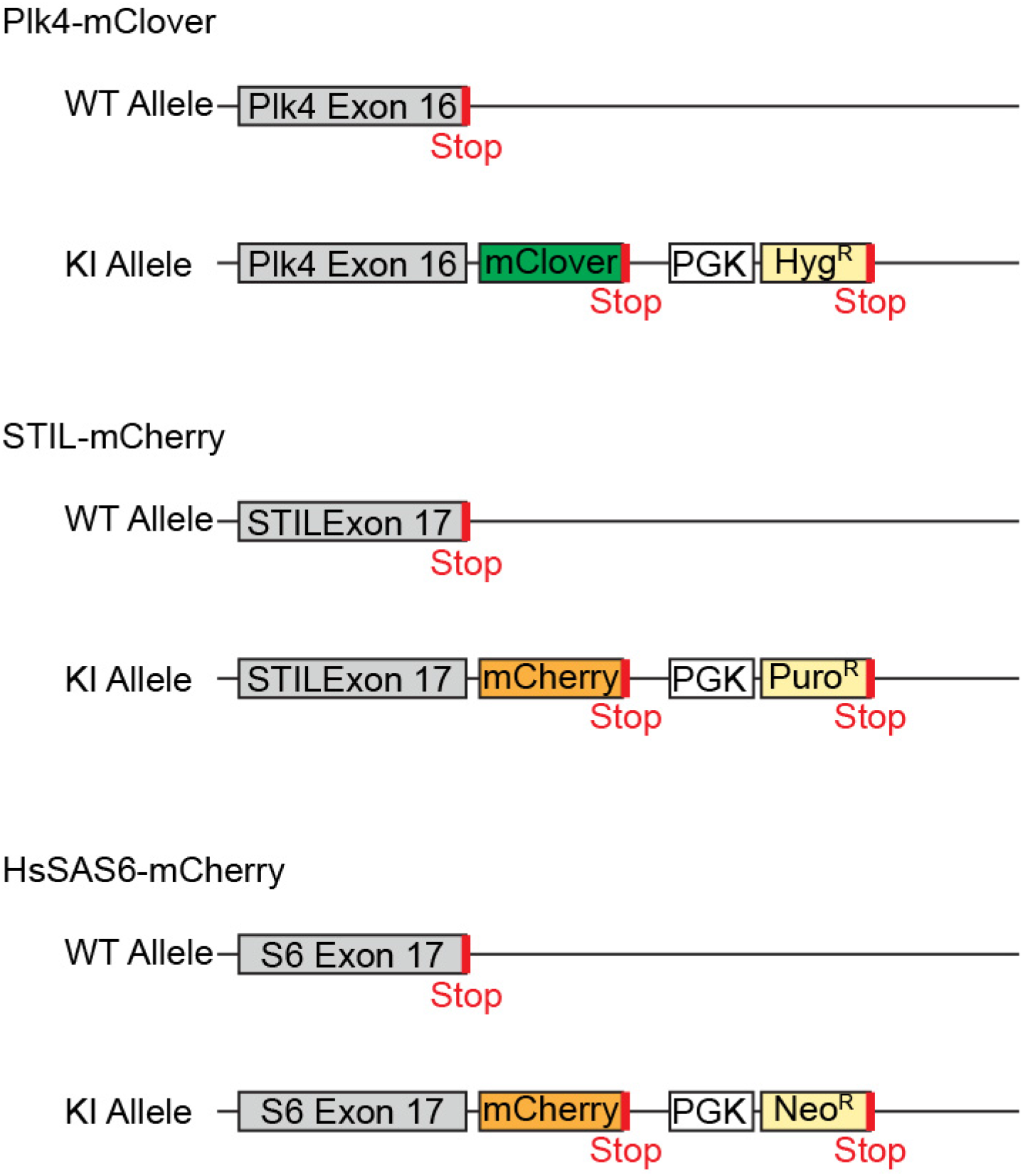
Endogenous tagging of the core components. The endogenous tagging system used for Plk4, STIL, and HsSAS6 are schematically shown.

## MATERIALS AND METHODS

### KEY RESOURCES

#### Antibodies

Plk4 (1:250; Merck, MABC544)

STIL (1:500; Abcam, ab89314)

HsSAS6 (1:500; Santa Cruz, sc-81431)

CEP152-N (1:1,000; Bethyl, A302-479A)

Pericentrin (1:1,000, Abcam, ab4448 or ab28144)

#### siRNAs

STIL (ThermoFisher, Silencer Select #s12863)

HsSAS6 (ThermoFisher, Silencer Select #s46487)

Negative control (ThermoFisher, Silencer Select #4390843)

#### Chemicals

Centrinone (MedChem Express, HY-18682).

## EXPERIMENTAL MODEL AND SUBJECT DETAILS

### Cell lines

HCT116 cells were cultured in McCoy’s 5A medium (GE Healthcare) supplied with 10% FBS, 1% glutamine, and 1% penicillin/streptomycin. HeLa cells were cultured in DMEM medium supplied with 10% FBS and 1% penicillin/streptomycin. HCT116 cells were used throughout this study unless otherwise stated. For Plk4 inhibition, centrinone was added to the medium to a final concentration of 200 nM.

Cells were transfected with siRNA using Lipofectamine RNAiMAX, according to the manufacturer’s instructions, for 24 h (live-cell imaging) or 48 h (immunofluorescence) before commencing live-cell imaging or fixation.

The HCT116 cell lines in which target proteins were fluorescently tagged were produced by CRISPR-Cas9 genome editing. The mClover or mCherry sequence was inserted into the 3’-region of the Plk4, STIL, or HsSAS6 gene as shown in Figure S6, and cell clones were selected using hygromycin, puromycin, or neomycin (Natsume et al., 2016). For mClover-tagged HsSAS6, a protein domain called AID (auxin-induced degron (Natsume et al., 2016)) was inserted between HsSAS6 and mClover, because the cell line was originally made for other purposes. Cloned cells were genotyped using PCR, and the proper localization of the expressed proteins with the fluorescent tag was verified by immunofluorescence. For the HCT116 Plk4-mClover + STIL-mCherry cell line, we failed to obtain a cloned cell in multiple trials for unknown reasons; we therefore analyzed fluorescence-positive cells from bulk culture instead. The STIL-mCherry and HsSAS6-mClover cell lines were biallelic. For the rest of the cell lines, we were only able to obtain monoallelic cell lines, which we used in the study.

## METHOD DETAILS

### Immunofluorescence

Cells cultured on coverslips were fixed using cold methanol at −20 °C for 5 min. The fixed cells were washed three times with PBS and incubated in blocking buffer (1% BSA and 0.05% Triton X-100 in PBS) for 20 min at room temperature (RT). The cells were then incubated with primary antibodies in blocking buffer at 4 °C overnight, washed three times with PBS, and incubated with secondary antibodies in blocking buffer for 1 h at RT. The cells were stained with Hoechst 33258 (DOJINDO) in PBS for 5 min at RT, washed three times with PBS, and subsequently mounted with ProLong Gold (Thermo Fisher Scientific, #P36930).

### Microscopy

For general observations, we used an upright epifluorescence microscope (Zeiss Axio Imager 2) with a 100× oil-immersion objective (N.A. 1.4) and an AxioCam HRm camera or an inverted confocal microscope (Leica TCS SP8) equipped with a 63× oil-immersion objective (N.A. 1.4). Z-stacked confocal images were obtained at 0.13 μm intervals. Huygens Essential image processing software was used for image deconvolution.

For live-cell imaging, we used a spinning disc-based confocal microscope (Yokogawa, CV1000) equipped with a 60× oil-immersion objective (N.A. 1.35), a back-illuminated EMCCD camera, and a stage incubator supplied with 5% CO2. Typically, 20–40 fields of view were recorded every 10 min for up to 30 h in a single experiment, and each field contained 25 z-slices at 1.3 μm intervals, subsequently max-projected using ImageJ software. ImageJ software was also used for image analyses. For siRNA treatment, cells were transfected with siRNA 24 h prior to the commencement of imaging. For centrinone treatment, image acquisition was temporarily paused for the addition of centrinone. The imaging chamber was quickly returned and the procedure proceeded in the same way and at the same fields of view as before.

### Mathematical modeling

To simulate the time course of the components involved in centriole duplication throughout the cell cycle, we constructed a mathematical model as follows. The quantities of the components were defined as in Table S1.

**Table S1.** Notation used for the quantity of each component in the model

| Notation | $P4$ | $P4p$ | $STIL$ | $S6$ | $S6cw$ | $P4_{cyto}$ | $STIL_{cyto}$ | $S6_{cyto}$ |
| --- | --- | --- | --- | --- | --- | --- | --- | --- |
| Definition | Plk4 | Phosphorylated<br>Plk4 | STIL | HsSAS6<br>free<br>dimer | HsSAS6<br>in<br>cartwheel | Plk4 | STIL | HsSAS6 |
|  | Centriolar |  |  |  |  | Cytosolic |  |  |

As schematically summarized in Figure 4A, the time course of the quantity of the centriolar components at time *t* is expressed using the following ordinary differential equations:

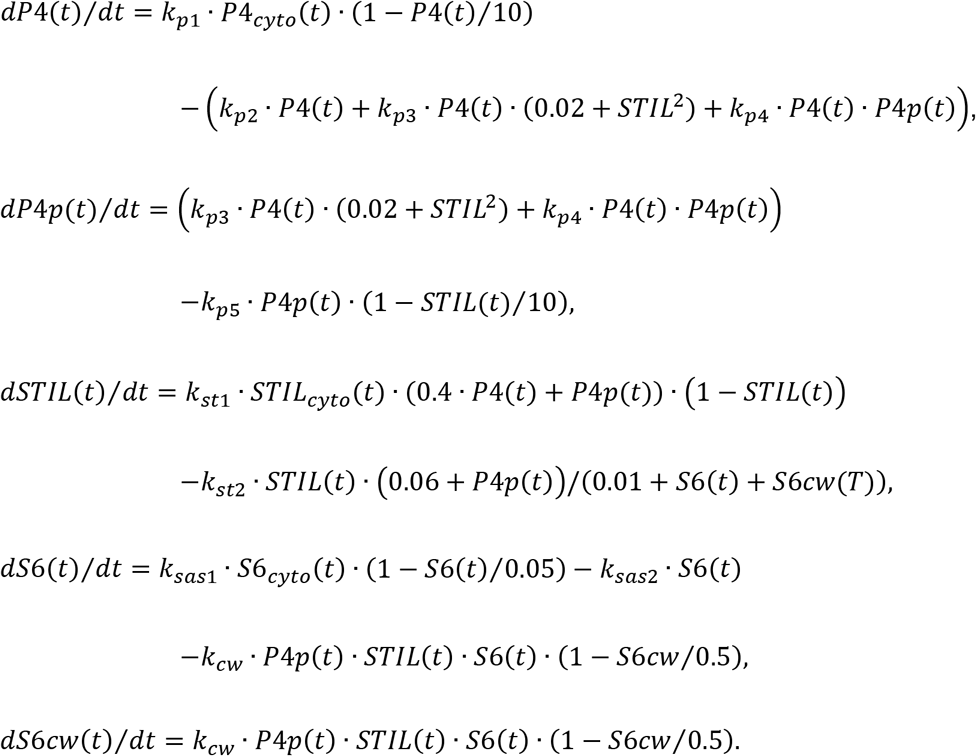

Note that all parameters, including quantity and time, are relative and dimensionless. For simplicity, STIL is assumed to be immediately phosphorylated by Plk4, so the non-phosphorylated and phosphorylated forms of STIL are not distinguished in the model. The time course of the expression levels of each component (i.e., the cytosolic fraction) are expressed as:

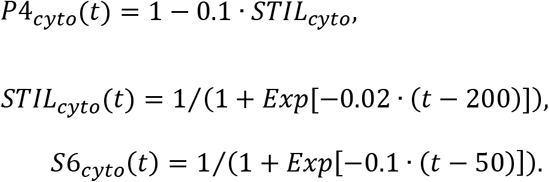

The initial quantity values were all set as 0. The parameter set used for the physiological condition is given in Table S2.

**Table S2.** Basic parameter settings for the physiological condition

| $k_{p1}$ | $k_{p2}$ | $k_{p3}$ | $k_{p4}$ | $k_{p5}$ | $k_{st1}$ | $k_{st2}$ | $k_{sas1}$ | $k_{sas2}$ | $k_{cw}$ |
| --- | --- | --- | --- | --- | --- | --- | --- | --- | --- |
| 0.02 | 0.02 | 0.2 | 0.02 | 0.2 | 0.16 | 0.08 | 0.01 | 0.01 | 2 |

Numerical solutions of the simultaneous differential equations were obtained using our original Mathematica program. For simulations of STIL- or HsSAS6-depletion (Figure 4D), the expression level of STIL or HsSAS6 was reduced to 10% and 1%, respectively, of the physiological level.

## QUANTIFICATION AND STATISTICAL ANALYSIS

All quantifications and statistical analyses were performed using ImageJ, Mathematica, Jupyter Notebook, and Microsoft Excel software. For live-cell imaging, the fluorescence intensity of regions of interest of the same size was measured using ImageJ on max projection images, and the fluorescence intensity of a no-cell region was used for background subtraction. For single-cell analyses (Figures 2A, 2B, S3, and S4), fluorescence was normalized to the maximum intensity of Plk4-mClover over the time course for each cell, and the moving average of fluorescence through time (± 3 time points) was calculated in Excel, for smoothing. The ring-filling indices of Plk4 were determined from oval profile plots with 64 sampling points, as described previously (Takao et al., 2018). The box-and-whisker plots (Figures 2D and 3C) and the parametric plots (Figures 2A, 2B, 4C, S3, and S4) were generated using our original code written in Python. We developed a Mathematica program to obtain cross-correlation functions (Figure 2D); the original data were zero-padded for the calculations.

